# Mapping Active-Site Conformational Ensembles Along Competing Catalytic Pathways of the Hairpin Ribozyme

**DOI:** 10.64898/2026.03.12.711346

**Authors:** Sélène Forget, Guillaume Stirnemann

## Abstract

The catalytic mechanism of the hairpin ribozyme has remained controversial for more than two decades, with different experimental approaches often supporting distinct mechanistic interpretations. In this work, we investigate the conformational landscape of the active site along several proposed reaction pathways using all-atom molecular dynamics simulations in explicit solvent combined with enhanced sampling techniques. Specifically, we employ Hamiltonian replica exchange simulations to extensively explore active-site conformations without relying on predefined collective variables, enabling a broad characterization of the structural ensembles associated with multiple protonation states along three candidate reaction mechanisms. Our simulations suggest that a dianionic general acid/general base pathway involving direct participation of A38 and G8 is unlikely to proceed through well-defined intermediates with catalytically competent geometries. In particular, states associated with G8 deprotonation and subsequent O2’ deprotonation exhibit strongly distorted active-site arrangements that appear poorly suited for reaction progression. Although highly synchronous proton-transfer steps cannot be excluded, the required deprotonation of G8 remains difficult to reconcile with neutral pH conditions. In contrast, monoanionic pathways in which the non-bridging oxygens of the scissile phosphate act as transient proton relays produce intermediates that sample geometries favorable for the nucleophilic addition and leaving-group elimination steps of the reaction. These mechanisms do not require direct catalytic involvement of G8 while remaining compatible with potential acid catalysis by protonated A38^+^ and a possible structural role of G8. Our results provide a unified conformational perspective on competing mechanistic scenarios. The ensembles generated here offer a foundation for future QM/MM and ML/MM calculations aimed at quantitatively resolving the free-energy landscapes governing hairpin ribozyme catalysis. Finally, the present strategy could easily be applied to other biomolecular systems with high conformational plasticity, including other ribozymes.

## Introduction

The discovery of catalytic RNAs, or ribozymes,^1–3^ fundamentally transformed our understanding of biological catalysis by demonstrating that chemical reactivity is not the exclusive domain of proteins. Known ribozymes typically catalyze either the ligation of two RNA fragments or their cleavage. RNA ligation in the absence of protein enzymes is considered essential for the emergence of the (putative) first RNA-based auto-catalytic systems.^4,5^ Self-catalyzed RNA cleavage remains highly relevant in extant biology, where it plays roles, for example, in viral RNA replication and RNA splicing.^6^ Although ribozymes catalyze a far narrower range of chemistries than protein enzymes, their catalytic mechanisms have been studied in depth;^5,7–9^ while a broadly consistent framework has emerged, significant open questions remain — well illustrated by the system examined here, the hairpin ribozyme (HpR).

The HpR accomplishes splicing through self-cleavage, that is, by replacing the 3’-5’ phosphodiester linkage between A-1 and G+1 with a 2’-3’ cyclic phosphate terminating linkage at the A-1 site,^10,11^ as illustrated in Figure 1. Although the HpR is not the only self-cleaving ribozyme, it is one of the few that can catalyze this reaction in the absence of divalent metal ions.^12–16^

**Figure 1:**
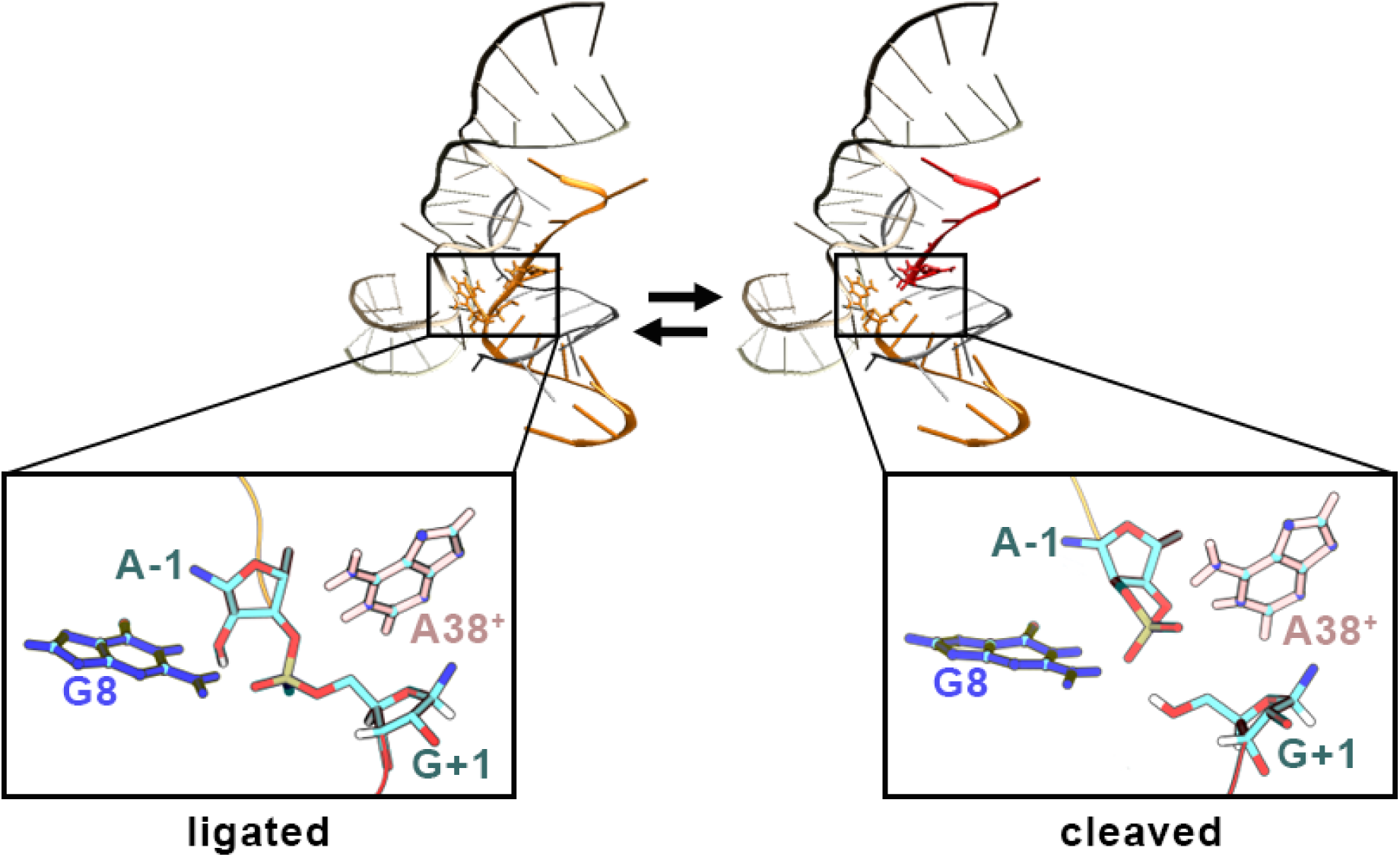
Hairpin ribozyme structures with a zoom on their catalytic sites, illustrating the self-cleavage/ligation reaction (left: ligated state, PDB ID: 2OUE ; right: cleaved state, PDB ID: 2P7D). The active site consists of the scissile bond between A-1 and G+1 residues, flanked by the nucleobases of G8 (blue) and A38^+^ (in pink).

The central step of this transphosphoesterification process is an addition–elimination reaction involving the formation of a pentavalent phosphorus transition state, which requires precise in-line alignment of the nucleophilic group (O2’), the central phosphorus atom, and the leaving group (O5’). From this framework, two key mechanistic questions arise: (i) how is the nucleophilic group activated prior to the addition step, and (ii) how is the leaving group activated prior to the elimination step? Although the critical catalytic roles of several HpR residues are well established, important questions remain regarding the precise nature of their involvement in the reaction mechanism, and these questions are similar for most other ribozymes, which catalyze the same chemical reaction.

Mutagenesis experiments,^17,18^ supported by high-resolution structural studies,^19–22^ have demonstrated that two conserved nucleobases, G8 and A38, are essential for catalytic activity and are located in close proximity to the reactive center.

On this basis, an initial mechanistic model was proposed.^23^ In the so-called dianionic mechanism, both G8 and A38 are directly involved: G8 acts as the general base, activating the O2’ nucleophile through proton abstraction (Figure 2, **D**_1_ → **D**_2_), while A38 serves as the general acid, reprotonating the O5’ leaving group following the elimination step (not shown in Figure 2).^23^ Although this model is consistent with pH-dependence experiments^24,25^ and with inferred transition-state geometries,^26^ it suffers from a major limitation: it requires prior deprotonation of the N1 atom of G8 (pKa*>*10),^27^ a condition that is unlikely to be met at physiological pH (Figure 2, R → **D**_1_).

**Figure 2:**
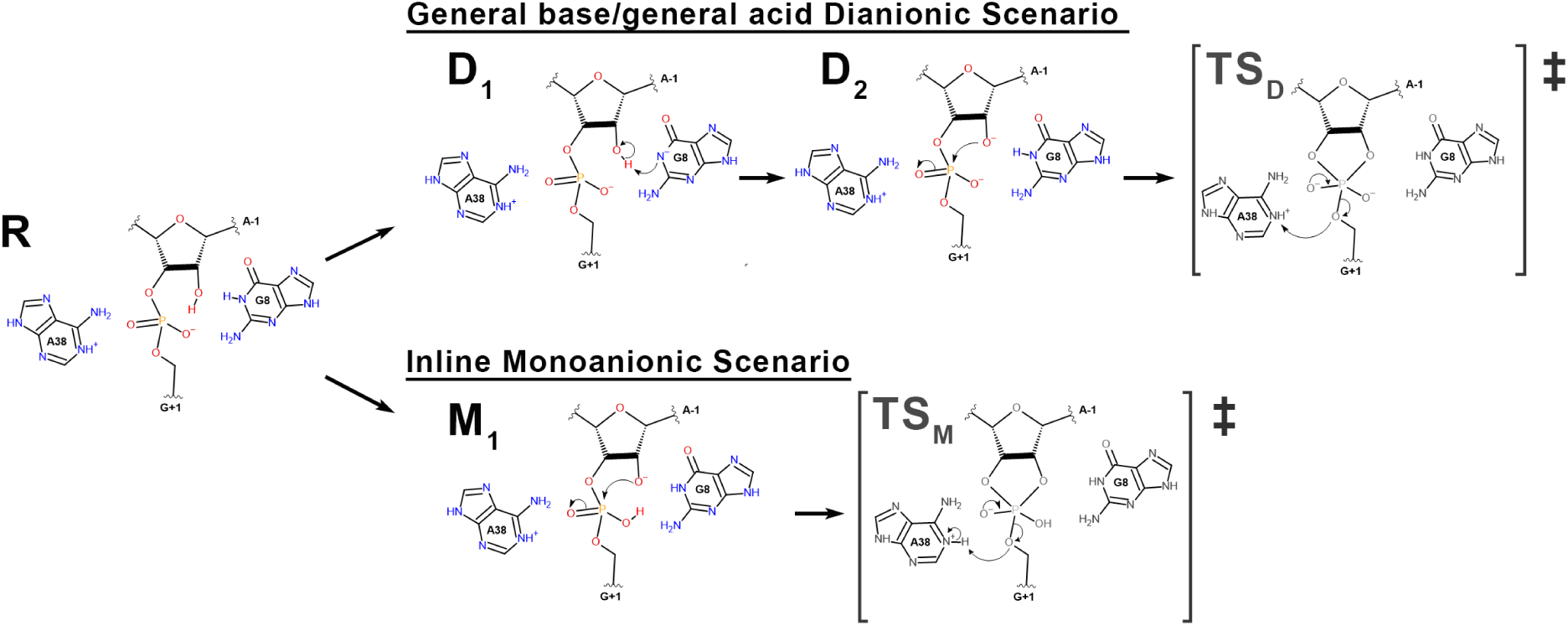
Proposed chemical mechanisms for the HpR self-cleavage reaction, from the reactant (R) to the respective transition states: the general acid-base mechanism, upper route (R → **D**_1_ → **D**_2_ → TS_D_), and the inline mono-anionic mechanism, lower route (R → **M**_1_ → TS_M_). Only the colored states are discussed in this study.

To resolve this inconsistency, an alternative mechanistic model has been proposed:^28^ the in-line mono-anionic mechanism (Figure 2, R → M_1_ → TS_M_). In this scenario, the nucleobases A38 and G8 play a critical role by contributing to the electrostatic stabilization of the transition state through a network of hydrogen bonds with atoms at the scissile junction. These non-covalent interactions pre-organize the catalytic site in a geometry favorable for reaction initiation. While the nucleobases remain properly positioned, deprotonation of the O2’ atom of A-1 is carried out by one of the non-bridging phosphate oxygens (NBO), resulting in the formation of a monoprotonated phosphate intermediate, **M**_1_. However, this mechanism is not easily connected to the experimentally observed pH dependence of the reaction, as argued by Kath-Schorr et al.^25^

The debate remains unresolved because the HpR is intrinsically self-reactive and cannot be structurally captured in its native, reactive state. Instead, studies rely on mimics of the HpR in which chemical activity is inhibited (either by removing the 2’-oxygen nucleophile, methylating it, or substituting other chemical groups with non-reactive analogs) to probe the catalytic site.^19,20^ Consequently, the catalytic site observed in these structures is distorted by the chemical modifications,^29^ and arguments based solely on crystal structures of such mimics, together with non-discriminative pH profiles, have not yielded a definitive mechanistic conclusion.

Complementary to experimental approaches, the HpR has been extensively studied in silico. Molecular dynamics (MD) simulations provide a molecular-level perspective that is not limited by the experimental necessity of chemical modifications. These computational studies can be divided into two main approaches. First, extended classical MD simulations of the full HpR have explored its conformational landscape using established force fields.^30–32^ Second, hybrid quantum–mechanical/molecular-mechanical (QM/MM) simulations have directly probed the catalytic site by modeling the chemical reaction steps. ^33–36^

These approaches are typically conducted sequentially, as QM/MM simulations require relevant pre-catalytic initial structures that are most reliably obtained from prior extensive classical MD sampling.

The conclusions of classical MD studies can be broadly grouped into two contrasting perspectives, depending on the assumptions made *a priori*. The first narrative arises from studies in which several catalytic site elements were explicitly constrained to enforce the formation of a specific hydrogen-bond network, consistent with the general dianionic scenario: G8:N1···A-1:O2’, A38:N1···G+1:O5’, G8:N2···G+1:O_pro_*_−_*_Sp_, and A38:N6···G+1:O_pro_*_−_*_Rp_.^31,37^

Although this local conformation appeared relatively stable over 500 ns of simulation, we previously demonstrated that its stability (labeled L_1_, visible in Figure 3) does not persist under extensive sampling.^38^ Instead, another local arrangement, L_2_ (Figure 3), represents the more stable conformation of the hairpin ribozyme in its pre-catalytic state. Both conformations are distinct from the one captured by the crystallographic resolution experiment,^19,20^ L_C_ (Figure 3).

**Figure 3:**
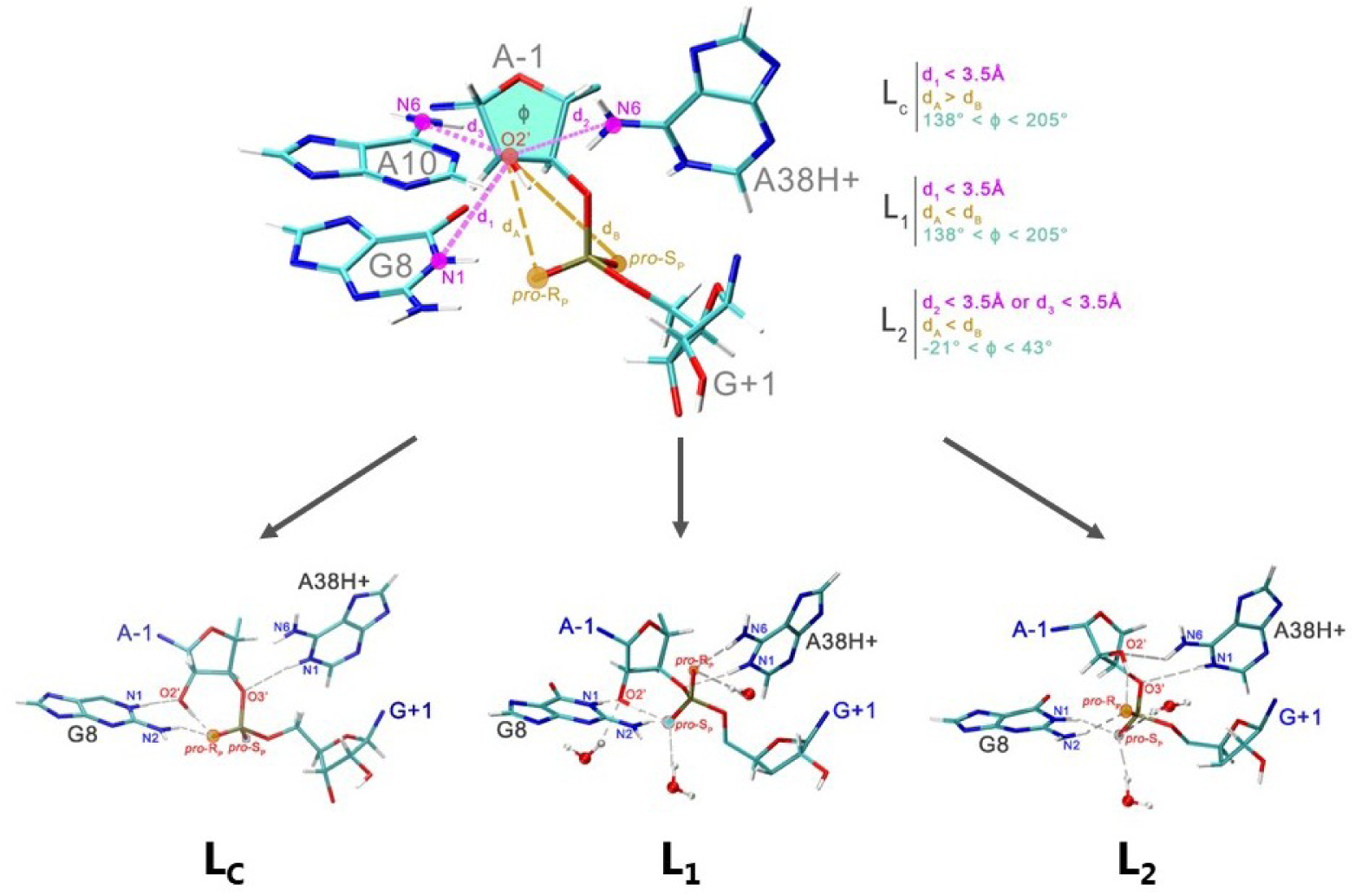
Key observables used to classify catalytic site conformations into L_C_, L_1_, L_2_ with details of their combinations as classification criteria. Representative structures of each conformation are shown below for the pre-catalytic protonation state. Note that a significant fraction of conformations cannot be attributed to any of these three basins, and are labeled ”others” whenever relevant.

These observations align more closely with the second narrative, which supports the inline monoanionic mechanism. In the L_2_ arrangement, G8 is protonated and positioned far from O2’, while O2’ forms a hydrogen bond with the non-bridging O_pro_*_−_*_Rp_ oxygen, placing the catalytic site in an ideal geometry for this mechanistic pathway. Furthermore, Kumar et al. explored the conformational landscape following transfer of the O2’ proton to O_pro_*_−_*_Rp_ and demonstrated that the catalytic site subsequently adopts an arrangement optimal for the addition–elimination steps, with an O2’–P–O5’ in-line attack angle (IAA) exceeding 140°.^32^ Complementary to classical MD simulations, several studies have employed QM/MM simulations to investigate the chemical activity of the HpR catalytic site. This approach allows explicit calculation of the electronic structure within the active site and enables evaluation of the (free) energy barriers separating reactants from products. In principle, such analyses can identify the mechanistic pathway that is most energetically favorable.^33,35,39^ Overall, QM/MM studies indicate that the monoanionic and dianionic mechanisms exhibit comparable free-energy barriers, suggesting that both pathways are energetically feasible and may either coexist or compete during HpR catalysis.

Far from resolving the debate, these in silico studies have instead underscored the complexity of the catalytic site’s conformational landscape and the difficulty of drawing definitive conclusions, given the inherent limitations of classical MD (force-field accuracy and sampling) and the computationally demanding, limited sampling accessible in QM/MM simulations.

Building on our previous studies, which demonstrated the critical importance of extensive conformational sampling of the ribozyme pre-catalytic state, ^29,38^ we aim to complement the existing in silico investigations of the HpR by specifically probing intermediate structures along plausible reaction pathways. Although such studies have been suggested before,^30,32^ none has applied the systematic, comprehensive sampling we propose here.

This strategy provides valuable mechanistic insights by evaluating the feasibility of individual steps at each stage of the reaction. Moreover, these simulations yield equilibrated structures along the different routes, providing the foundation required for subsequent QM calculations.

We find that the pre-catalytic site of the HpR arranges coherently with the in-line monoanionic mechanism, adopting flat IAAs only after the proton has been transferred to one of the two non-bridging phosphate oxygens, thus positioning the site favorably for the subsequent addition–elimination step. This behavior is observed for both the pro-Sp and pro-Rp pathways, suggesting the existence of two possible in-line monoanionic sub-scenarios. In contrast, our simulations suggest that the general acid–base mechanism is very unlikely to proceed in a stepwise manner. First, deprotonation of G8, which is required for this nucleobase to act as a general base, causes it to move away from the scissile junction. Second, proton transfer from O2’ to G8 generates a negative charge on O2’ that appears to repel the phosphate group, thereby preventing nucleophilic attack. Unless these steps occur in a strongly concerted fashion, they constitute a sequence of highly improbable chemical events.

## Methods

### Simulation parameters and Analysis

All simulations were performed using GROMACS 2022.3^40^ patched with PLUMED 2.7.^41,42^ The visualization and analysis of the REST2 trajectories were performed using the VMD software,^43^ with the Collective Variables Dashboard,^44^ and several custom scripts (available on the GitHub repository of the group) built on both the MDAnalysis^45,46^ and the Barnaba^47^ Python libraries.

### Forcefields

Consistent with the conclusions drawn from our previous work, ^38^ we employed the Amber14 ff99bsc0*χ*_OL3_*ɛζ*_OL1_ force field^48^ for RNA, the TIP3P water model, and Joung-Cheatham parameters^49^ for K^+^ and Cl*^−^* ions.

This force field lacks parameters for several non-standard residues in our HpR intermediates (all listed in Table S1), which we added as follows. Parameters for N1-protonated A38 (RAP), N1-deprotonated G8 in the **D**_1_ intermediate (RGM), and the joined A-1/G+1 residues forming monoprotic phosphate species (GAA with protonated O_pro_*_−_*_Rp_; RAA with deprotonated A-1:O2’*^−^*) were taken from the literature^38^ and manually adapted. Parameters for the O_pro_*_−_*_Sp_-protonated monoprotic phosphate (RSA + GSA) were derived from RAA and GAA by inverting the charges on O_pro_*_−_*_Rp_ and O_pro_*_−_*_Sp_ while preserving all other charge distributions. The O2’-deprotonated A-1*^−^* residue in the **D**_2_ dianionic intermediate (AD) was parametrized using Gaussian^50^ and AmberTools^51^ to obtain RESP charges and generate appropriate atom types. All parameters can be found in the Supporting Information.

### Initial configurations

Most simulations were initiated from the crystallographic structure of the minimal HpR pre-catalytic state (PDB ID: 2OUE), ^19^ after manual removal of the methyl group from the O2’ atom of A-1, following the protocol used in our previous work.^38^ The only exceptions are the initial structures for the in-line monoanionic intermediates through pro-Sp that were initiated from an L^pucker^ frame extracted from the trajectory provided by Mlynsky et al.^31,38^ Indeed, the corresponding phosphate conformation is very far from the crystal structure reference, and would naturally follow the occurrence of the L^pucker^ conformation where the corresponding proton transfer would occur, as detailed later in the results section. With our enhanced sampling scheme, it is expected that the conformational sampling would be little sensitive to the initial starting structure upon proper equilibration, as shown before for the pre-catalytic state. ^38^

### Simulation parameters

Electrostatic interactions were calculated using the particle-mesh Ewald method^52^ with a Verlet cutoff scheme and 10.0 Å cutoff for Lennard-Jones interactions. The PME grid spacing was set to 0.12 nm using cubic interpolation. All bonds involving hydrogen atoms were constrained using LINCS,^53^ and a 2 fs time step was used.

### System set up and equilibration

All systems were solvated in TIP3P water using the GROMACS solvate command. Crystallographic waters were retained, while crystallographic ions were replaced with 0.2 M KCl. All solvated ribozymes underwent minimization, equilibration, and production runs following the protocol in our previous work . ^38^ Briefly, the solvent was progressively equilibrated via 5000-steps energy minimization, followed by 500 ps of NVT and NPT equilibration at T = 298.15 K and P=1 bar, all while maintaining the solute with 1000 kJ/mol/nm² positional restraints on the nucleic backbone. The solvent being equilibrated, the solute restraints were gradually reduced through three successive relaxation stages (1000, 100, and 10 kJ mol*^−^*^1^ nm*^−^*^2^). Finally, the system was equilibrated for 100 ps in NVT and 100 ps in the NPT ensemble.

### Replica Exchange with Solute Tempering (REST2)

All HpR simulations were performed using replica exchange with solute tempering (REST2),^54,55^ using parameters identical to our previous work: ^38^ we employed 24 replicas with rescaling factors ranging from 1 to 0.667, exchange attempts every 1 ps at 300 K, at 300 K and 500 ns production runs per replica. Exchange acceptance rates ranged from 13–20%.

### Descriptors

We characterized catalytic site conformations in ligated states using the labeling scheme from our previous work^38^ (Figure 3). This scheme distinguishes three main conformations based on A-1 sugar pucker, O2’ hydrogen bonding, and phosphate orientation. In L_1_, the A-1 residue is in the C2’-endo puckering pseudo-rotation angle, the O2’ atom is involved in a H-bond with G8:N1, and the phosphate is oriented such that the O2’ is closer to the O_Sp_ than the O_Rp_ oxygen. In L_2_, A-1 adopts C3’-endo pucker, O2’ hydrogen bonds to A38:N6 or A10:N6, and O2’ is closer to O_pro_*_−_*_Rp_. In L_C_, A-1 adopts C2’-endo pucker, O2’ hydrogen bonds to G8:N1, and O2’ is closer to O_pro_*_−_*_Rp_. Frames that do not meet these criteria are classified as ”others”.

### Convergence

The convergence of the simulations of all ligated intermediates was assessed both at the global scale, using the eRMSD metric^56^ over time, and at the local scale, using the fraction of frames in each conformational category (see above) per time block. Figure S1 A–E details the time evolution of these metrics for all systems, with converged time ranges indicated.

### Clustering

Each REST2 trajectory was clustered based on inter-nucleobase distances (specifically, the distances between the heavy atoms at the extremities of the Watson-Crick edges: N3 and O4 on uracils, N1 and N6 on adenines, N2 and O6 on guanines, and O2 and N4 on cytosines). To reduce dimensionality and only keep relevant distances, we selected atom pairs within 5 Å in 5–95% of converged trajectory frames. We used the YACARE algorithm,^57^ a clustering approach specifically designed to address the challenges of analyzing replica exchange simulations: it reorders the frames and detects clusters as density centers within the conformational similarity space, allowing for noise classification. The key observables are plotted along the reordered frames in Figure S2 and Figure S4, revealing the size of the clusters and of the noise. Snapshots of the representative structures of certain clusters are displayed underneath to visualize their catalytic site conformations.

## Results

### Strategy

We performed all-atom molecular dynamics simulations using Hamiltonian replica exchange (HREX) in explicit solvent, employing the Amber14 ff99bsc0*χ*_OL3_*ɛζ*_OL1_ RNA force field with TIP3P water, as described in the Methods section. This choice of force field and sampling strategy, which is a critical aspect of molecular simulations of RNA,^58,59^ is motivated by our previous studies where we compared several parametrizations.^29,38^ In particular, among the many force fields we tested, this model yields consistent and reliable results for the pre-catalytic **R** state. We have also shown that conventional brute-force MD simulations fail to achieve convergence on microsecond timescales, whereas HREX provides substantially improved sampling of the ribozyme conformational space while requiring a tractable number of replicas compared to traditional temperature replica exchange. For this system, the design of specific and relevant collective variables (CVs) is particularly challenging, if not impractical, rendering CV-biased approaches unsuitable.

In the following, we successfully study the different states along the possible reaction pathways (Figure 2). With the exception of the pre-catalytic state, studied in detail in our previous study, the current work focuses on the intermediate states along possible reaction pathways. Note that we will refer to these states as *intermediates*, despite the absence of evidence for their metastability or existence (therefore, this term does not correspond to any assumption about their chemical relevance). We will only assume that they correspond to different, successive protonation/deprotonation events and assess the relative stability and conformations of the corresponding states.

### Mono-Anionic Scenario

We start this conformational exploration of mechanistic intermediates with the mono-anionic mechanism, which begins with a proton transfer from the O2’ to one of the two non-bridging oxygens of the scissile phosphate, leading to the formation of the **M**_1_ intermediate, sometimes called the ”AP” intermediate^32^ (see Figure 2, R→**M**_1_). From there, the deprotonated O2’ oxygen attacks the phosphorus atom via an addition-elimination mechanism, forming a pentavalent phosphorus transition state (TS_M_ in Figure 2), and then eliminating the O5’ of the following residue which deprotonates the NBO of phosphate that was protonated upon M_1_ formation.

Because it involves the protonation of a phosphate NBO, the mono-anionic mechanism can proceed via two different oxygens, namely the O_pro_*_−_*_Sp_ (O1P) and the O_pro_*_−_*_Rp_ (O2P) pathways (Figure 3). Although they appear symmetrical on paper, these two pathways are not equivalent in terms of local arrangements and energy. Indeed, they imply the at least transient formation of an H-bond between HO2’ and the NBO proton acceptor, meaning that they are initiated from distinct conformations of the catalytic site. In other words, the catalytic site conformational ensemble that is observed for the pre-catalytic R state can inform on the most likely proton acceptor. Should O2’ be involved in an H-bond with O_pro_*_−_*_Rp_, the proton transfer would occur towards this NBO and the **M**_1_ intermediate would correspond to a protonated O_pro_*_−_*_Rp_ species. Likewise, if O2’ is rather facing O_pro_*_−_*_Sp_, the following mono-anionic scenario will likely present an **M**_1_ protonated O_pro_*_−_*_Sp_. We thus examine the conformations of the active site in its pre-catalytic state.

### Pre-catalytic state

The extensive exploration of the conformational landscape of the pre-catalytic HpR has been the subject of previous studies.^31,38^ We observed that the pre-catalytic site arranges in two major conformations, L_2_ and L_1_, which differ in their H-bond patterns (see Methods and Figure 3). As we have previously shown, L_2_ is the most stable conformation and L_1_ is metastable, i.e., it can be observed in simulations with lifetimes exceeding hundreds of nanoseconds but always falls into the L_2_ basin upon sufficient sampling,^38^ for all of the many investigated forcefields, including the one used in the present study. The crystallographic conformation L_c_, which corresponds to a structure containing a 2’-O-methylation, is never observed in the simulations.^29,38^

In L_2_, there is an H-bond between A-1:O2’···G+1:O_pro_*_−_*_Rp_, while in L_1_, an A-1:O2’···G+1:O_pro_*_−_*_Sp_ H-bond is present (see Figure 3). As shown in Figure 4, the conformational exploration provided by the HREX simulations shows that L_2_ (in cyan) is very dominant over the converged part of the pre-catalytic system simulations (81.4% of the frames). In this state, HO2’ always remains closer to G+1:O_pro_*_−_*_Rp_ than to O_pro_*_−_*_Sp_ and is ready to be exchanged between the A-1:O2’ and G+1:O_pro_*_−_*_Rp_, which would suggest that the O_pro_*_−_*_Rp_ pathway would naturally follow from this conformation.

**Figure 4:**
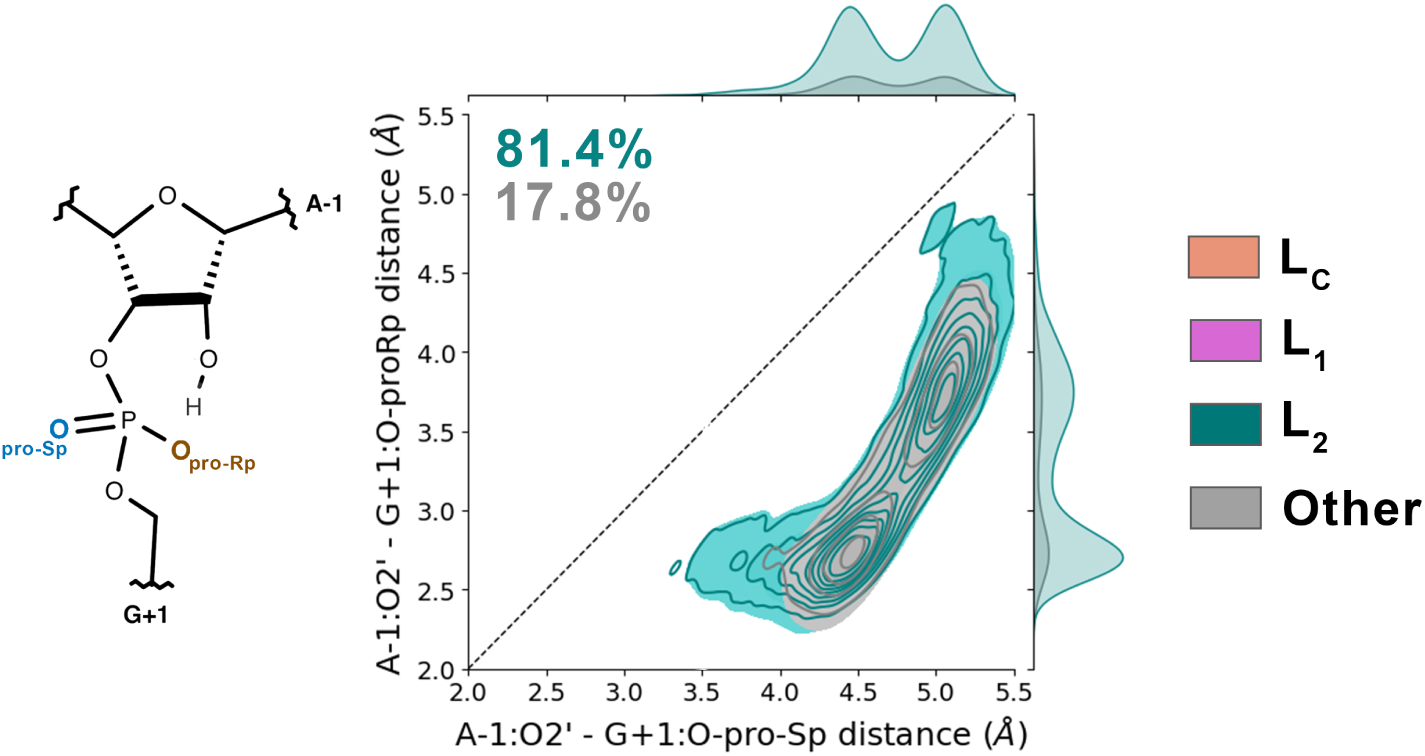
2D distribution of A-1:O2’···G+1:O_pro_*_−_*_Sp_ (O1P) versus A-1:O2’···G+1:O_pro_*_−_*_Rp_ (O2P) distances for the converged portion of the pre-catalytic HpR simulation (R in Figure 2). The inset shows a sketch of the scissile bond with non-bridging phosphate oxygens O_pro_*_−_*_Sp_ and O_pro_*_−_*_Rp_ color-coded. Percentages of frames in each labeled conformation are shown in the upper left (see Methods and Figure 3 for conformational labeling). The dashed diagonal line (x = y) indicates equal distances to both NBOs, to help identify which oxygen is closer to O2’.

On the contrary, the L_1_ conformation of the pre-catalytic state exhibits an H-bond between A-1:O2’···G+1:O_pro_*_−_*_Sp_, a natural precursor of a proton transfer towards O_pro_*_−_*_Sp_. The L_2_ → L_1_ conformational transition therefore requires crossing a free-energy barrier whose magnitude remains to be evaluated, but we cannot exclude that the proton transfer could occur via a metastable conformation, the formation of which would be an intermediate step in the overall mechanistic pathway. Without this debate being settled, we now choose to consider both O_pro_*_−_*_Sp_ and O_pro_*_−_*_Rp_ accordingly intermediate states that follow the proton transfer.

### AP intermediates: proton transfer through O_pro*−*Rp_

Starting with the O_pro_*_−_*_Rp_ pathway, we first discuss the situation in which A-1:O2’ is deprotonated and the O_pro_*_−_*_Rp_ of the scissile phosphate is protonated (Figure 5B). Following deprotonation, the negatively charged O2’*^−^* nucleophile is expected to attack the phosphorus atom. The key observables for this event are the IAA and d_O2_*′−_−_*_P_, which we represent in a 2D distribution plot in Figure 5B. A local minimum, absent in the pre-catalytic state 2D distribution, emerges in the IAA’155° and d_O2_*′−_−_*_P_’3.25 Å region. This conformation is ideally arranged for nucleophilic attack and represents 35.1% of the converged REST2 simulation.

**Figure 5:**
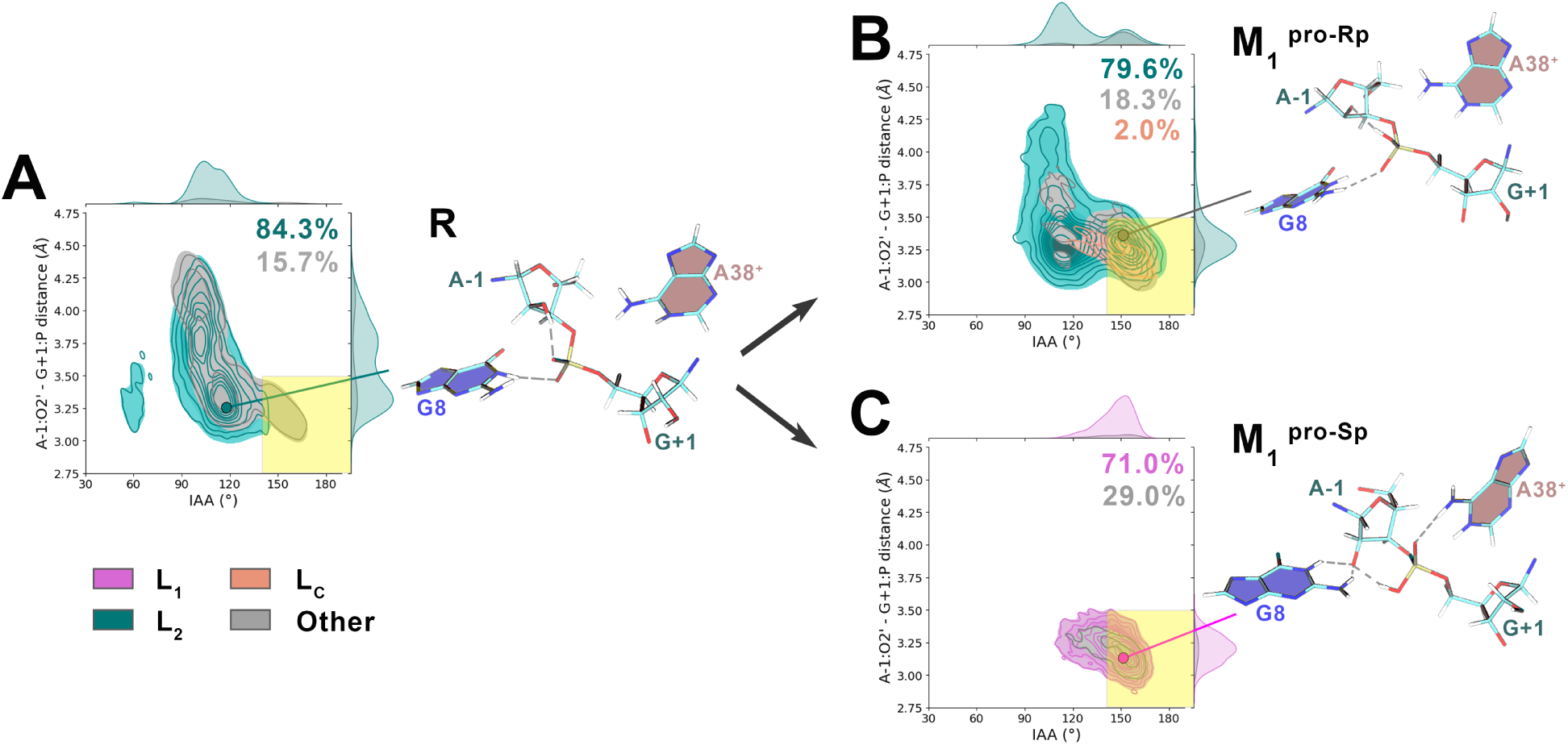
2D distributions of the A-1:O2’···G+1:P distance (d_O2′−P_) and inline attack angle (IAA) for the converged portions of REST2 simulations of monoanionic HpR intermediates: (A) pre-catalytic state, (B) O_pro_*_−_*_Rp_-protonated intermediate, and (C) O_pro_*_−_*_Sp_-protonated intermediate. Distributions are colored by conformation type (see Methods and Figure 3). Insets show representative structures from the dominant high-IAA, low-d_O2′−P_ clusters (cluster 0 for A, cluster 4 for B, cluster 0 for C). See Methods and Figure 3 for conformational labeling and clustering methodology, and Figure S2 for detailed clustering results. The zone labeled as ”reactive” (d_O2′−P_≤3.5Å & IAA≥140°) is highlighted in yellow.

These high IAA conformations correlate with the loss of G+1:O_pro_*_−_*_Rp_···G8:N2 and A-1:O2’*^−^*···A38:N6 H-bonds and the formation of a G+1:O_pro_*_−_*_Sp_···G8:N2 and, more remarkably, the G+1:O_pro_*_−_*_Rp_···G+1:N2 H-bonds (clusters 2, 3, 4 visible in Figure S2A). Now protonated, O_pro_*_−_*_Rp_ oxygen is poised to act as proton donor and finds a partner in the 2-amino group of G+1, which adopts a geometry at the boundary between the syn and trans conformations, like in all other investigated protonation states and the reference crystal structure.^20^ As the O_pro_*_−_*_Rp_ NBO and N2 atom approach each other, the IAA increases, reaching values above 140° when the H-bond forms (see Figure S3).

These results suggest that G+1 itself may play a structural role in stabilizing the catalytic site geometry for the monoanionic mechanism. To our knowledge, this specific involvement of G+1 has never been discussed in previous computational studies of the HpR. However, this hypothesis finds support in experimental mutagenesis studies, ^60^ which revealed a clear correlation between the internal equilibrium towards ligation and the nature of the +1 residue: the greater the number of hydrogen bond donors/acceptors on the nucleobase, the higher the ligation rate, consistent with a direct structural role for G+1 in organizing the catalytic site.

### AP intermediates: proton transfer through O_pro*−*Sp_

Similarly to the pro-Rp pathway, the protonated O_pro_*_−_*_Sp_ intermediate constitutes the state from which the nucleophilic attack occurs, and its catalytic viability can be assessed through analysis of the IAA and d_O2_*′−_−_*_P_ distributions. Figure 5C shows the corresponding 2D plot, and reveals that the catalytic site of this intermediate is ideally arranged for the nucleophilic attack, with 73% of the frames in the reactive region (IAA *>* 140° and d_O2_*′−_−_*_P_ *<* 3.5 Å). The representative structure of the main cluster (Figure 5C, or cluster 0 in Figure S2B) displays a typical arrangement: the transferred proton forms a strong H-bond between A-1:O2’*^−^* and G+1:O_pro_*_−_*_Sp_, while on the opposite side of the scissile phosphate, A38 nucleobase nitrogens form H-bonds with the O_pro_*_−_*_Rp_ and remain in close proximity to G+1:O5’, with 88% of frames showing d_O5_*′_−_*_A38:N1_ *<* 3.5 Å.

Overall, the catalytic site adopts a position favorable for nucleophilic attack and consistent with experimental interpretations: both G8 and A38 form key H-bonds that maintain the favorable inline arrangement, although they do not directly participate in the reaction itself. This arrangement is remarkably rigid, with our clustering analysis identifying only four global structure clusters, all sharing the L_1_ catalytic site conformation (Figure S2B). Quite noticeably, the active site geometry in that case recalls that of the previous studies that constrained these H-bonds in the ribozyme pre-catalytic state in order to achieve reactive arrangements seen in experimental structures of TS-mimics.^37^

### Dianionic Scenario

#### Deprotonation of G8

We begin examination of the dianionic scenario with the **D**_1_ intermediate. In principle, the mechanism should start with the deprotonation of G8:N1 (Figure 2, **R** → **D**_1_), yet most HpR studies overlook this initial mechanistic step. While we follow this convention, we emphasize that this step should not be taken for granted: the high pKa of G8:N1 (*>*10.5)^27^ renders deprotonation unlikely under physiological conditions, and no other plausible base close enough to G8 could catalyze this deprotonation.

From **D**_1_, the dianionic mechanism proceeds via proton transfer from A-1:O2’ to the deprotonated G8:N1*^−^* (**D**_1_ → **D**_2_). Previous studies reported significant catalytic site distortion with G8:N1*^−^*,^30^ which our simulations confirm. The G8*^−^* residue moves away from the reactive center, with the G8:N1*^−^*···A-1:O2’ distance averaging over 8 Å in the most stable configuration. The free energy profile along this distance (Figure 6) reveals that bringing G8:N1*^−^* close enough to form a hydrogen bond with A-1:O2’ (G8:N1*^−^*···A-1:HO2’ distances around 2–2.5 Å) requires overcoming a barrier that should largely exceed 3 kcal/mol. Yet, this substantial free energy cost merely positions the groups for potential hydrogen bonding, and the actual proton transfer would still be thermodynamically unfavorable given the large pKa difference between G8:N1 (pKa *>* 10) and typical alcohol groups (pKa ∼ 16-18).

**Figure 6:**
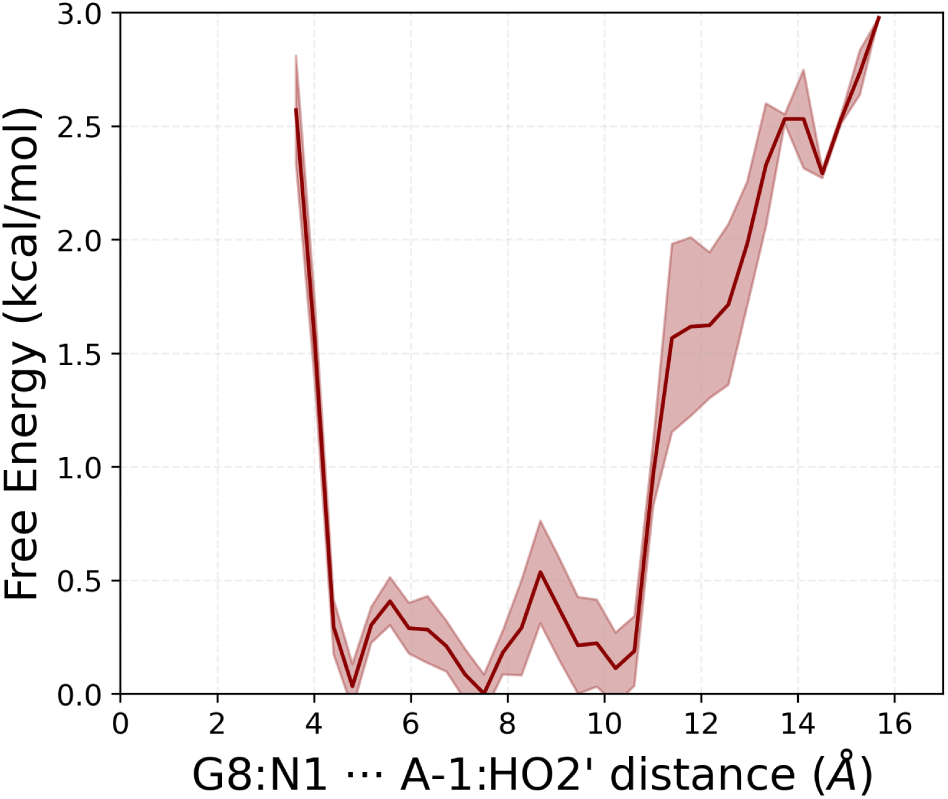
Free energy profile as a function of the A-1:O2’···G8:N1*^−^* distance for the D_1_ intermediate, averaged over three 100 ns time blocks, with calculated standard deviation shown as error bars (in transparency). According to the dianionic mechanism, deprotonated G8:N1*^−^* should abstract the O2’ proton to activate the nucleophile for the subsequent addition step.

We also observe that, upon deprotonation, G8:N1 becomes highly accessible to the surrounding water (Figure S5), implying that this state would necessarily be very transient: reprotonation by water would be far more favorable than deprotonation of the O2’.

The displacement of G8*^−^* occurs predominantly within an L_2_ arrangement (71.4% of converged frames, see Figure 7A), with most conformational features persisting, such as the A-1:O2’···A38:N6 H-bonding and the C3’-endo sugar pucker conformation of A-1 (Figure S4A). However, electrostatic repulsion between the negatively charged G8*^−^* and the phosphate group and the loss of the G8:N1H···phosphate H-bonds liberate the phosphate. The scissile junction can now rotate into multiple conformations and can even achieve flat IAAs. Yet, in all of these aligned conformations G8*^−^* remains too distant to abstract the O2’ proton and activate the O2’ nucleophile (clusters 10 and 11, Figure S4A).

**Figure 7:**
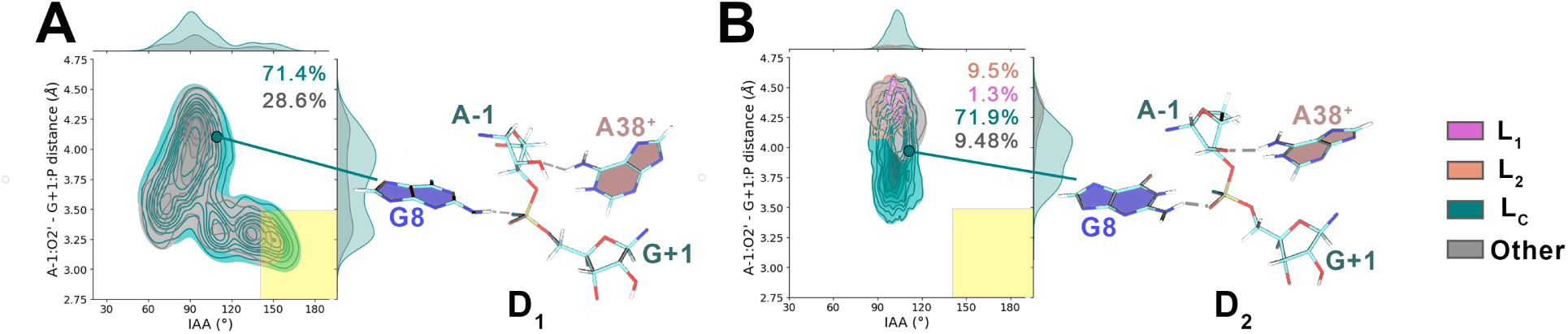
2D distributions of inline attack angle (IAA) versus A-1:O2’···G+1:P distance (d_O2′−P_) for the converged portions of REST2 simulations of dianionic HpR intermediates: (A) deprotonated G8*^−^* acts as general base and (B) nucleophile A-1:O2’*^−^* is deprotonated. Distributions are colored by conformation type. Insets show representative structures from the main clusters (cluster 0 for A, cluster 4 for B). See Methods and Figure 3 for conformational labeling and clustering methodology, and Figure S4 for detailed clustering results. The zone labeled as ”reactive” (d_O2′−P_≤3.5Å & IAA≥140°) is highlighted in yellow.

### Activation of O2’ with a basic G8

Following the hypothetical proton transfer (**D**_1_ → **D**_2_), the deprotonated O2’*^−^* nucleophile should attack the phosphorus via an addition-elimination mechanism. This requires an IAA ≥ 140° and a short A-1:O2’*^−^*···G+1:P distance. However, our simulations reveal that the catalytic site does not adopt geometries meeting these criteria. The IAA distribution shows a single peak with a maximum at 120°, while d_O2_*′−_−_*_P_ remains above 3.75 Å (Figure 7B).

Our clustering analysis reveals that here, since the O2’*^−^* is now negatively charged, it is itself repelled by the phosphate, and rather interacts with A38 and A10 amines when A-1*^−^* adopts C3’-endo pucker conformation (clusters 1, 2, 3, 4 and 7, 8, visible in Figure S4B), or with G8:N1 when A-1*^−^* adopts C2’-endo (clusters 0 and 5). None of these catalytic site arrangements is favorable for the nucleophilic attack.

## Discussion

We now discuss our results in the context of available experimental data. Experimental studies unequivocally support the conclusion that G8 and A38 are critical for hairpin ribozyme catalysis, as mutations at these positions have pronounced effects on catalytic activity. ^17,18^ However, the interpretation of pH-dependence data has been the subject of considerable debate.^17,23,25,61,62^ In particular, although the functional importance of A38 is universally acknowledged,^22,30,63,64^ the precise role of G8 is contentious. While some studies argue against its involvement as a general base,^65^ others attribute the lack of a clear pH-dependent signature to a high pKa that lies outside the experimentally accessible range but is shifted into this range upon mutation, consistent with a role as a general base.^25,61^ Support for a general acid–general base mechanism is further provided by structures of transition-state analogs obtained using chemical modifications, which clearly show A38 and G8 positioned appropriately to function as the general acid and general base, respectively.^65^

With a pKa of G8:N1 estimated to be above 10,^27^ only a very small fraction of the ribozyme population would occupy this state under equilibrium conditions at neutral pH. Moreover, the lifetime of such a state is expected to be extremely short, i.e., on the order of a few picoseconds, before reprotonation by surrounding water molecules occurs. Beyond these kinetic and thermodynamic considerations, the corresponding ribozyme state **D**_1_ exhibits a markedly distorted active site. In particular, G8 is displaced away from the reactive center as a result of electrostatic repulsion and gets exposed to the solvent, leading to a substantial reorganization of the hydrogen-bonding network relative to the pre-catalytic state. This disruption is unfavorable for catalysis because it compromises the precise positioning and stabilizing interactions required to promote nucleophilic attack. Consistent with this picture, our simulations indicate that shortening the A-1:O2’···G8:N1*^−^* distance incurs a significant free-energy penalty of at least several kcal/mol, to which must be added the cost of the proton transfer between these two groups, whose pKa difference is close to 8.

Although the presence of Mg^2+^ cation in the active site is a common strategy found in several ribozymes to reduce electrostatic repulsion and maintain negatively-charged residues close together, this is unlikely to be the case here. There is evidence that HpR is able to function without magnesium,^16^ and thiol substitution studies suggest that when present, it does not interact with the scissile phosphate.^12,14,15^

We next investigate the **D**_2_ conformation that follows this proton transfer. Likely because both the phosphate group oxygens and the O2’*^−^* are now negatively charged (the O2’*^−^* charge is parametrized at -0.82331), the active site fails to adopt geometries conducive to catalysis. In particular, the nucleophile, phosphorus atom, and leaving group are poorly aligned, and the nucleophile–phosphorus distances remain large. In addition, the ribose ring of the reacting residue preferentially adopts puckering conformations that differ from those observed in other states along the reaction pathway. Finally, the hydrogen-bond network is strongly perturbed and deviates substantially from the network generally considered critical within the framework of the general acid/general base mechanism. Polarizable force fields, mixed quantum/classical schemes, and machine-learned interatomic potentials could describe the local active-site geometry more accurately, especially in the presence of such highly localized, like-signed charges. Their cost, and the attendant limits on sampling, together with the need for systematic experimental validation, has so far restricted their use in large-scale RNA simulations.

Taken together, these results suggest that the general acid/general base mechanism, while attractive and consistent with several experimental observations, involves a sequence of molecular events that is unlikely given the geometries of the associated states and leads to intermediates that do not preserve an active-site structure favorable for catalysis. Nevertheless, these findings do not completely rule out this mechanism. If all steps were fully concerted and no metastable intermediates existed along the reaction pathway, these putative “intermediates” would not need to be thermodynamically relevant. It is noteworthy that QM/MM studies have described both sequential^39^ and concerted^33,34^ dianionic scenarios. In any case, this scenario does not resolve the fundamental issue of G8:N1 deprotonation (pKa’10.6^27^), which is highly improbable under neutral conditions and remains an open and often insufficiently addressed question.

For the monoanionic scenario involving phosphate-assisted proton transfers, the pre-catalytic state conformations are ideally arranged for proton transfer between O2’ and the O_pro_*_−_*_Rp_ oxygen, with a stable pre-existing hydrogen bond between these two groups that could readily precede such a transfer. Both M_1_ intermediates, bearing a proton on either the O_pro_*_−_*_Rp_ or O_pro_*_−_*_Sp_ oxygen and a deprotonated O2’*^−^*, are well positioned for the subsequent nucleophilic attack, displaying favorable in-line geometries and short A-1:O2’*^−^*···G+1:P distances.

In addition, the pro-Sp M_1_ intermediate exhibits an overall hydrogen-bonding network that closely matches what is generally expected to support progression along the reaction coordinate, including stabilizing interactions that preserve the relative positioning of key catalytic groups. This state adopts an L_1_ conformation, which has consistently been identified in previous studies as a reactive geometry. ^31,37^ The associated ribose puckering is also compatible with this assignment, remaining similar to that observed in other catalytically competent states along the pathway. Notably, the crystallographic structure of the HpR captured in a transition-state analog^66^ aligns closely with the hydrogen-bond network observed in the highly stable pro-Sp intermediate, in particular through the presence of the G8:N2···G+1:O_pro_*_−_*_Sp_ and A-1:O2’···G8:N1 H-bonds.

The main limitation of the monoanionic scenario is that it appears, at first glance, to be inconsistent with the experimentally observed pH dependence of activity, which increases at low pH before reaching a plateau. ^17^ This behavior suggests the involvement of a titratable group with a pKa close to 6.^23,25^ If a phosphate NBO were the sole titratable group directly participating in the reaction, one would instead expect an increase in activity only at very low pH values (approximately 1–2), followed by a plateau. However, invoking the participation of a phosphate NBO does not exclude the involvement of A38, which could plausibly account for the experimentally observed pH signature. There is also evidence from previous simulation studies that a canonical A38 significantly distorts the active site geometries,^30^ as confirmed by complementary simulations (see Figure S6). In this respect, just as the signature associated with G8 acting as a base lies outside the experimentally accessible pH window, the titration range of a phosphate NBO would fall at extremely low pH values that are likewise not experimentally relevant.

Evidence for the involvement of G8 as a general base largely relies on the observed decrease in catalytic activity at high pH values (8–9) when G8 is mutated to a residue with a lower pKa.^24^ However, this observation does not constitute decisive proof that G8 is directly involved in catalysis. First, as discussed above, G8 could play an indirect role, as is the case for many residues in protein enzymes, by contributing to the electronic stabilization of the active site. Such an effect could account for both the pH dependence of the catalytic rate^17,23,24,61,67^ and the sensitivity of the reaction to mutations at this position. ^17,62^

Second, the models commonly used to relate catalytic rates to the fraction of available acid and base species as a function of pH appear to be only qualitatively consistent with experimental data.^61^ In particular, the amplitude of the pH dependence of the rate is relatively modest, typically no more than a factor of three between optimal and minimal activity, compared to what would be expected based on these models. Such limited variations could equally well arise from subtle but functionally significant pH-induced changes in the overall ribozyme structure, a phenomenon that would not be surprising and that has been documented in related systems.^68,69^

In addition, it is noteworthy that the monoanionic scenario involving phosphate-assisted proton transfers closely resembles the uncatalyzed reaction mechanism in bulk water. This mechanism has recently been investigated by some of us using deep neural network–based interatomic potentials,^70,71^ as well as by others in earlier studies.^72,73^ In this framework, the role of the ribozyme active-site residues would primarily be to provide an electrostatic environment that lowers the free-energy barriers of the reaction,^74^ while the reaction itself would proceed along a pathway that is fundamentally similar to that observed in bulk solution. These studies suggested a barrier of 15–18 kcal/mol for the first proton transfer between the nucleophile and a non-bridging oxygen (NBO) of the phosphate in the uncatalyzed reaction in water,^70,71^ compared with around 11 kcal/mol for the same event in the ribozyme.^36^ These numbers should be compared with great care, as the methodologies differ markedly — machine-learned interatomic potentials in the first case, QM/MM calculations in the second — but it is notable that both studies used the same DFT functional (BLYP-D3) as reference data and converged on a similar mechanism, albeit with a lower barrier in the ribozyme.

## Conclusion

Using Hamiltonian replica exchange molecular dynamics, we have mapped the conformational landscape of the hairpin ribozyme active site along three putative reaction pathways. The molecular mechanism of catalysis by the hairpin ribozyme has been intensely debated over the past two decades, with different experimental groups often reaching conflicting conclusions.^25,61,63,65^ Each experimental technique comes with intrinsic limitations and typically provides indirect information about the mechanism. As a result, distinct mechanistic scenarios can sometimes be consistent with the same experimental data. Previous computational studies have contributed important molecular insights; however, they have not fully resolved the controversy, partly due to questions regarding the employed force fields, the extent of sampling, and the choice of starting structures.^30–32,37^ Moreover, mixed quantum–classical simulations have suggested that multiple mechanisms may compete, exhibiting very similar activation barriers.^33–36^

Our results support the view that the dianionic general acid/general base mechanism, while consistent with several experimental observations, faces severe structural obstacles: the key intermediates along this pathway adopt active-site geometries that are highly distorted and poorly compatible with catalytic progression, and the deprotonation of G8:N1 required to initiate it is thermodynamically improbable at neutral pH. The monoanionic pathways, in which the non-bridging phosphate oxygens act as proton relays, present a structurally more favorable picture: the pre-catalytic state is pre-organized for the initial proton transfer, and the resulting intermediates sample geometries broadly consistent with the requirements for nucleophilic attack. On this basis, we conclude that the currently available experimental and computational evidence is insufficient to unambiguously discriminate between the two leading mechanistic scenarios, and that the monoanionic pathways deserve greater consideration than they have so far received.

This conclusion is, however, subject to several key limitations. The first concerns the inherent constraints of non-polarizable force fields, particularly when describing intermediate states that involve non-canonical protonation patterns, which cannot be readily benchmarked against extensive experimental data. Although we addressed this issue in detail in previous work,^38^ where we compared multiple force fields for the ribozyme pre-catalytic state, we cannot exclude the possibility that alternative parameterizations or other levels of descriptions might yield different active-site geometries for certain intermediates. We note, however, that the force field employed here was selected based on its overall consistency with available experimental observations.

A second limitation concerns sampling. Although Hamiltonian replica exchange provides substantially improved coverage of conformational space compared to standard molecular dynamics, full convergence of the relevant ensembles on microsecond timescales is not guaranteed even with enhanced sampling. Recent advances in machine learning offer promising avenues to address this, including data-driven design of improved reaction coordinates^75^ and further enhancements to replica-exchange efficiency.^76,77^

A final issue is that the intermediates discussed throughout this study are, at best, metastable. If some are in fact short-lived transition-state-like species, detailed analysis of their active-site conformations may have limited discriminatory power between competing pathways.

Despite these limitations, the conformational ensembles generated here provide a unified and internally consistent view of how changes in protonation state induce active-site rearrangements along three mechanistic pathways. From an experimental standpoint, directly probing the involvement of the phosphate non-bridging oxygens as proton relays would be the most informative next step. From a computational perspective, we anticipate that these ensembles will serve as robust starting configurations for mixed quantum/classical and ML/MM calculations,^78^ which could address the transient nature of intermediates and the accuracy limitations of classical force fields. Ultimately, resolving the mechanistic debate will require free-energy landscapes for the competing pathways; the extensive conformational sampling presented here constitutes an essential foundation for that next step.

## Author contribution statement

S.F. and G.S. conceived the research. S.F. conducted the simulations, S.F. and G.S. analyzed the results. S.F. and G.S. wrote and reviewed the manuscript.

## Acknowledgments

The research leading to these results has received funding from the European Research Council under the European Union’s Eighth Framework Program (H2020/2014-2020)/ERC Grant Agreement No. 757111 (G.S.). The simulations presented here benefited from access to the HPC resources of TGCC under the allocation A0130811005 made by GENCI (Grand Equipement National de Calcul Intensif).

## Data availability statement

All simulation inputs and trajectory data used to generate figures are available in our Zenodo repository at 10.5281/zenodo.20511966. Analysis scripts are available at https://github.com/stirnemann-group/RNA-analysis. A tutorial with related protocols is available from the PLUMED-tutorials website. ^79^ All simulations were performed with standard open-source software as described in the Methods.

## Supporting Information

## Simulation parameters

The Amber14 ff99bsc0*χ*_OL3_*ɛζ*_OL1_ force field is available at https://kfc.upol.cz/ff_ol/. Non-standard residues not included by default in this force field are listed in Table S1, with parameter details adapted from previous studies provided below.

## RGM: guanine residue deprotonated on its N1 atom (rtp format), adapted from ^1^

~~~
[ RGM ]
[ atoms ]
P P 1.166200 1
O1P O2 -0.776000 2
O2P O2 -0.776000 3
O5’ OS -0.498900 4
C5’ CI 0.055800 5
H5’1 H1 0.067900 6
H5’2 H1 0.067900 7
C4’ CT 0.106500 8
H4’ H1 0.117400 9
O4’ OS -0.354800 10
C1’ CT -0.022700 11
H1’ H2 0.200600 12
N9 N* 0.000500 13
C8 CP 0.012500 14
H8 H5 0.133900 15
N7 NB -0.552100 16
C5 CB -0.030100 17
C6 C 0.770700 18
O6 O -0.703400 19
N1 NC -0.874400 20
C2 CA 0.914300 21
N2 N2 -0.961000 22
H21 H 0.357300 23
H22 H 0.357300 24
N3 NC -0.820800 25
C4 CB 0.333400 26
C3’ C7 0.202200 27
H3’ H1 0.061500 28
C2’ CT 0.067000 29
H2’1 H1 0.097200 30
O2’ OH -0.613900 31
HO’2 HO 0.418600 32
O3’ OS -0.524600 33
~~~

## RAA: adenine residue deprotonated on its sugar O2’ atom (rtp format), adapted from^2^

~~~
[ RAA ]
[ atoms ]
P P 1.166200 1
O1P O2 -0.776000 2
O2P O2 -0.776000 3
O5’ OS -0.498900 4
C5’ CI 0.055800 5
H5’1 H1 0.067900 6
H5’2 H1 0.067900 7
C4’ CT 0.106500 8
H4’ H1 0.117400 9
O4’ OS -0.354800 10
C1’ CT 0.039400 11
H1’ H2 0.200700 12
N9 N* -0.025100 13
C8 C5 0.200600 14
H8 H5 0.155300 15
N7 NB -0.607300 16
C5 CB 0.051500 17
C6 CA 0.700900 18
N6 N2 -0.901900 19
H61 H 0.411500 20
H62 H 0.411500 21
N1 NC -0.761500 22
C2 CQ 0.587500 23
H2 H5 0.047300 24
N3 NC -0.699700 25
C4 CB 0.305300 26
C3’ C7 0.202200 27
H3’ H1 0.061500 28
C2’ CT 0.067000 29
H2’1 H1 0.097200 30
O2a O2 -0.776000 31
O3’ OS -0.524600 32
~~~

## GSA: monoprotic guanine residue with the O_pro_*_−_*_Sp_ protonated (rtp format), adapted from^2^

~~~
[ GSA ]
[ atoms ]
P P 1.166200 1
O1n OH -0.613900 2
O2P O2 -0.776000 3
HOP HX 0.418600 4
O5’ OS -0.498900 5
C5’ CI 0.055800 6
H5’1 H1 0.067900 7
H5’2 H1 0.067900 8
C4’ CT 0.106500 9
H4’ H1 0.117400 10
O4’ OS -0.354800 11
C1’ CT 0.019100 12
H1’ H2 0.200600 13
N9 N* 0.049200 14
C8 CP 0.137400 15
H8 H5 0.164000 16
N7 NB -0.570900 17
C5 CB 0.174400 18
C6 C 0.477000 19
O6 O -0.559700 20
N1 NA -0.478700 21
H1 H 0.342400 22
C2 CA 0.765700 23
N2 N2 -0.967200 24
H21 H 0.436400 25
H22 H 0.436400 26
N3 NC -0.632300 27
C4 CB 0.122200 28
C3’ C7 0.202200 29
H3’ H1 0.061500 30
C2’ CT 0.067000 31
H2’1 H1 0.097200 32
O2’ OH -0.613900 33
HO’2 HO 0.418600 34
O3’ OS -0.524600 35
~~~

## GAA: monoprotic guanine residue with the O_pro_*_−_*_Rp_ protonated (rtp format), adapted from^2^

~~~
[ GAA ]
[ atoms ]
P P 1.166200 1
O1P O2 -0.776000 2
O2n OH -0.613900 3
HOP HX 0.418600 4
O5’ OS -0.498900 5
C5’ CI 0.055800 6
H5’1 H1 0.067900 7
H5’2 H1 0.067900 8
C4’ CT 0.106500 9
H4’ H1 0.117400 10
O4’ OS -0.354800 11
C1’ CT 0.019100 12
H1’ H2 0.200600 13
N9 N* 0.049200 14
C8 CP 0.137400 15
H8 H5 0.164000 16
N7 NB -0.570900 17
C5 CB 0.174400 18
C6 C 0.477000 19
O6 O -0.559700 20
N1 NA -0.478700 21
H1 H 0.342400 22
C2 CA 0.765700 23
N2 N2 -0.967200 24
H21 H 0.436400 25
H22 H 0.436400 26
N3 NC -0.632300 27
C4 CB 0.122200 28
C3’ C7 0.202200 29
H3’ H1 0.061500 30
C2’ CT 0.067000 31
H2’1 H1 0.097200 32
O2’ OH -0.613900 33
HO’2 HO 0.418600 34
O3’ OS -0.524600 35
~~~

---

**Table S1:**
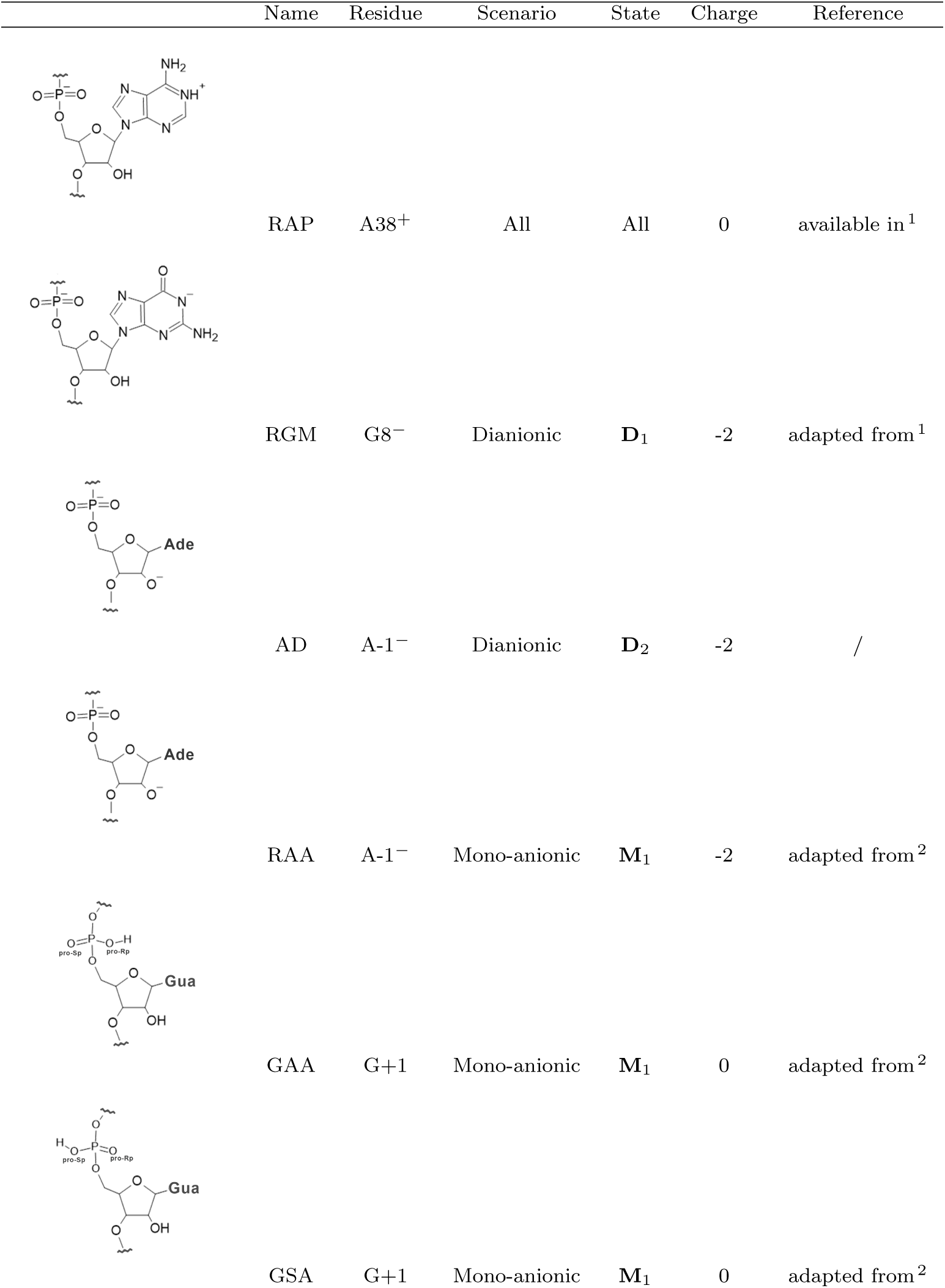
Summary of the non-standard residues of this study.

|  | Name | Residue | Scenario | State | Charge | Reference |
| --- | --- | --- | --- | --- | --- | --- |
|  | RAP | A38<sup>+</sup> | All | All | 0 | available in <sup>1</sup> |
|  | RGM | G8<sup>-</sup> | Dianionic | **D**<sub>1</sub> | -2 | adapted from <sup>1</sup> |
|  | AD | A-1<sup>-</sup> | Dianionic | **D**<sub>2</sub> | -2 | / |
|  | RAA | A-1<sup>-</sup> | Mono-anionic | **M**<sub>1</sub> | -2 | adapted from <sup>2</sup> |
|  | GAA | G+1 | Mono-anionic | **M**<sub>1</sub> | 0 | adapted from <sup>2</sup> |
|  | GSA | G+1 | Mono-anionic | **M**<sub>1</sub> | 0 | adapted from <sup>2</sup> |

**Figure S1:**
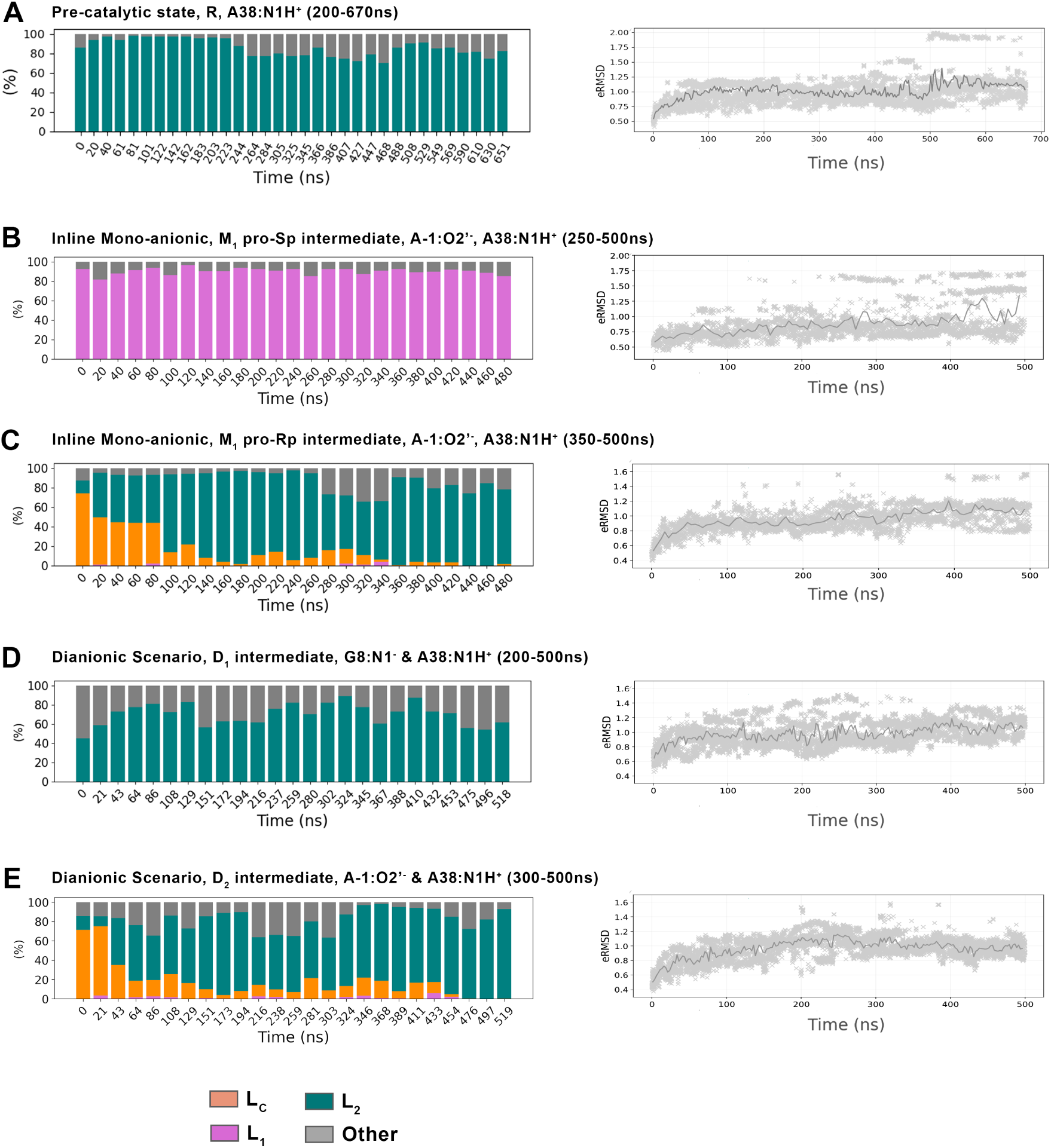
Time series of observables from the unscaled replica used to assess convergence. Left: local catalytic site behavior monitored through the fraction of frames in each labeled conformation per 20 ns time block (see Methods and Figure 3 for classification details). Right: global system behavior monitored through the eRMSD^3^ relative to each system’s initial structure. Convergence times determined from both metrics are indicated in parentheses.

**Figure S2:**
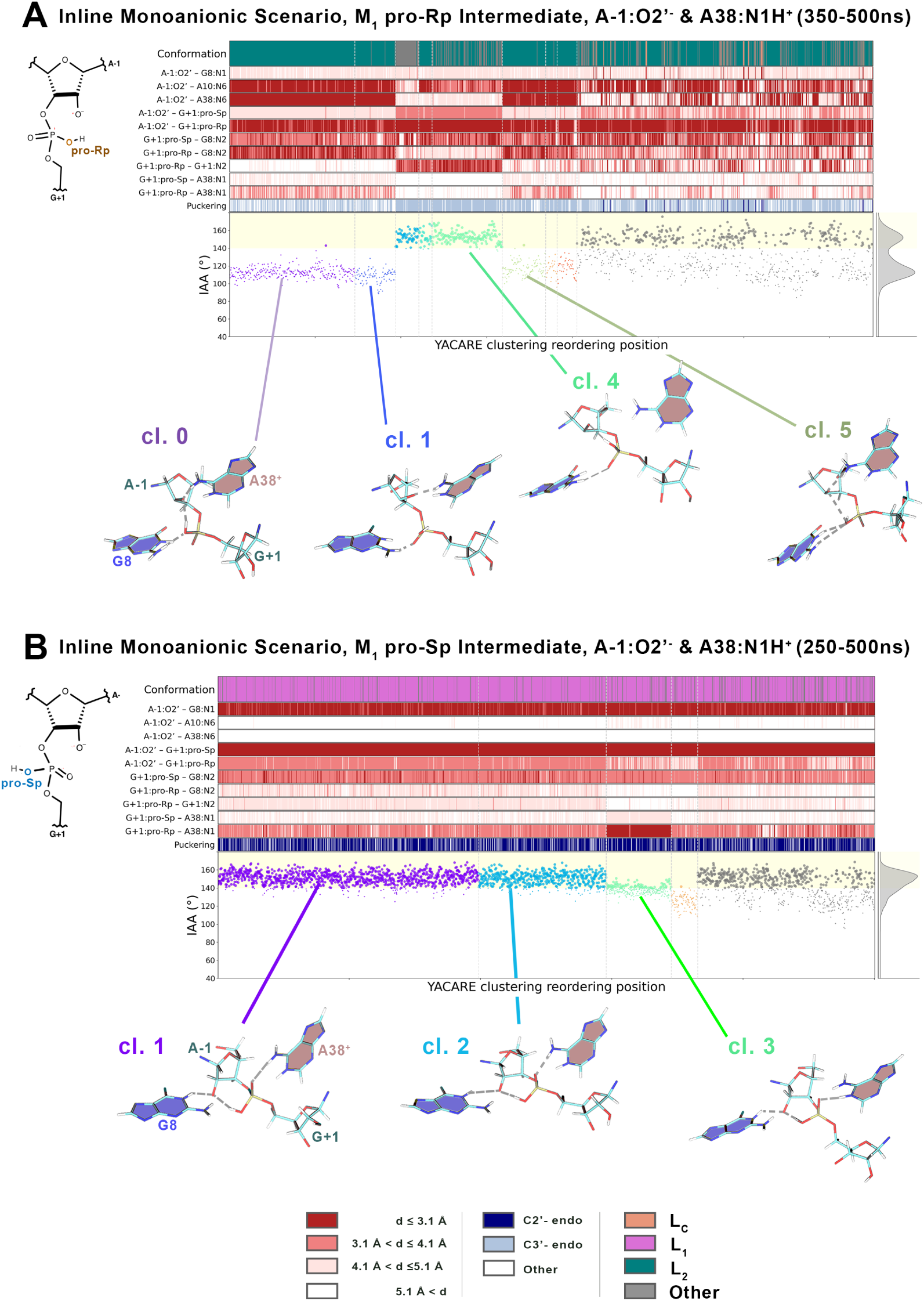
Evolution of key catalytic site observables for monoanionic intermediates, with frames ordered by clustering analysis: (A) O_pro_*_−_*_Rp_ protonated and (B) O_pro_*_−_*_Sp_ protonated M_1_ intermediate (see Figure 2). Row 1: conformational label (see Methods and Figure 3 for conformational labeling strategy); rows 2–10: heatmap of the key catalytic site distances; row 11: A-1 puckering pseudo-rotation conformations; row 12: inline attack angle (IAA) values. Below: representative structures from major clusters with hydrogen bonds (represented by dashed lines).

**Figure S3:**
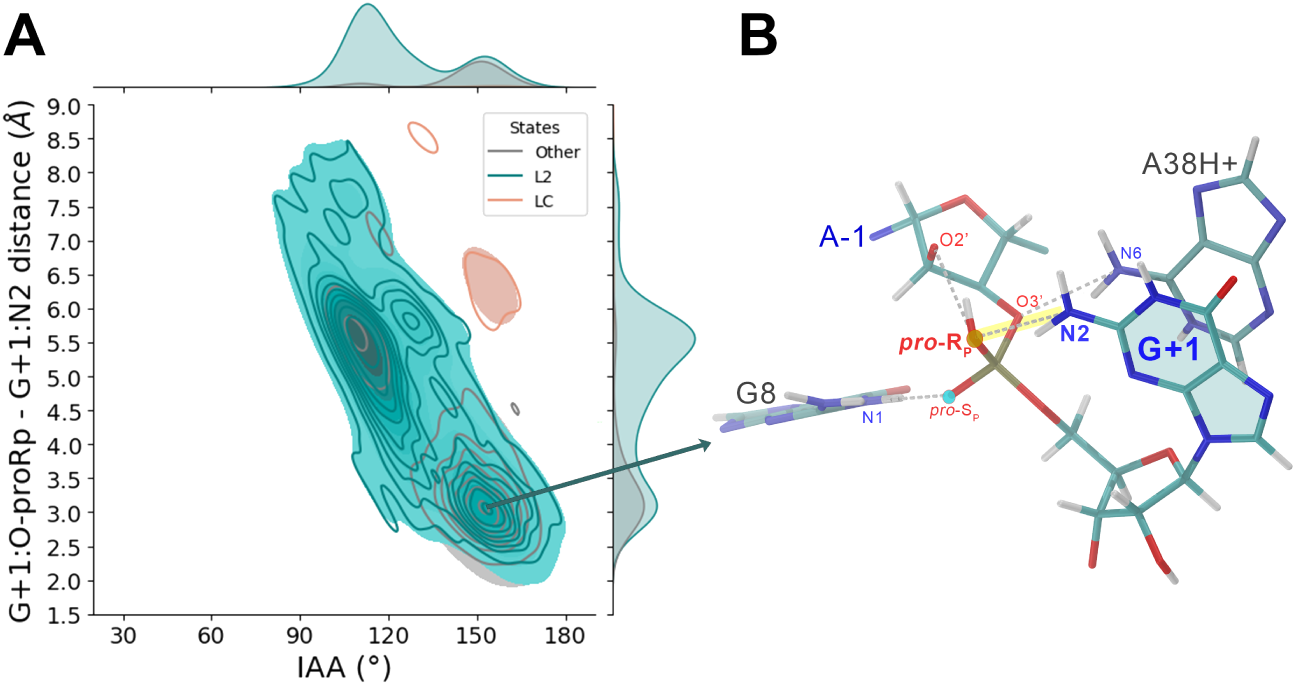
(A) 2D distribution of inline attack angle (IAA) versus G+1:O_pro_*_−_*_Rp_···G+1:N2 distance for the converged portions of REST2 simulations of monoanionic protonated O_pro_*_−_*_Rp_ HpR intermediates, colored by conformational label (see Methods and Figure 3 for conformational labeling strategy). (B) View of the catalytic site in an aligned conformation, with the H-bonds apparent in dashed lines.

**Figure S4:**
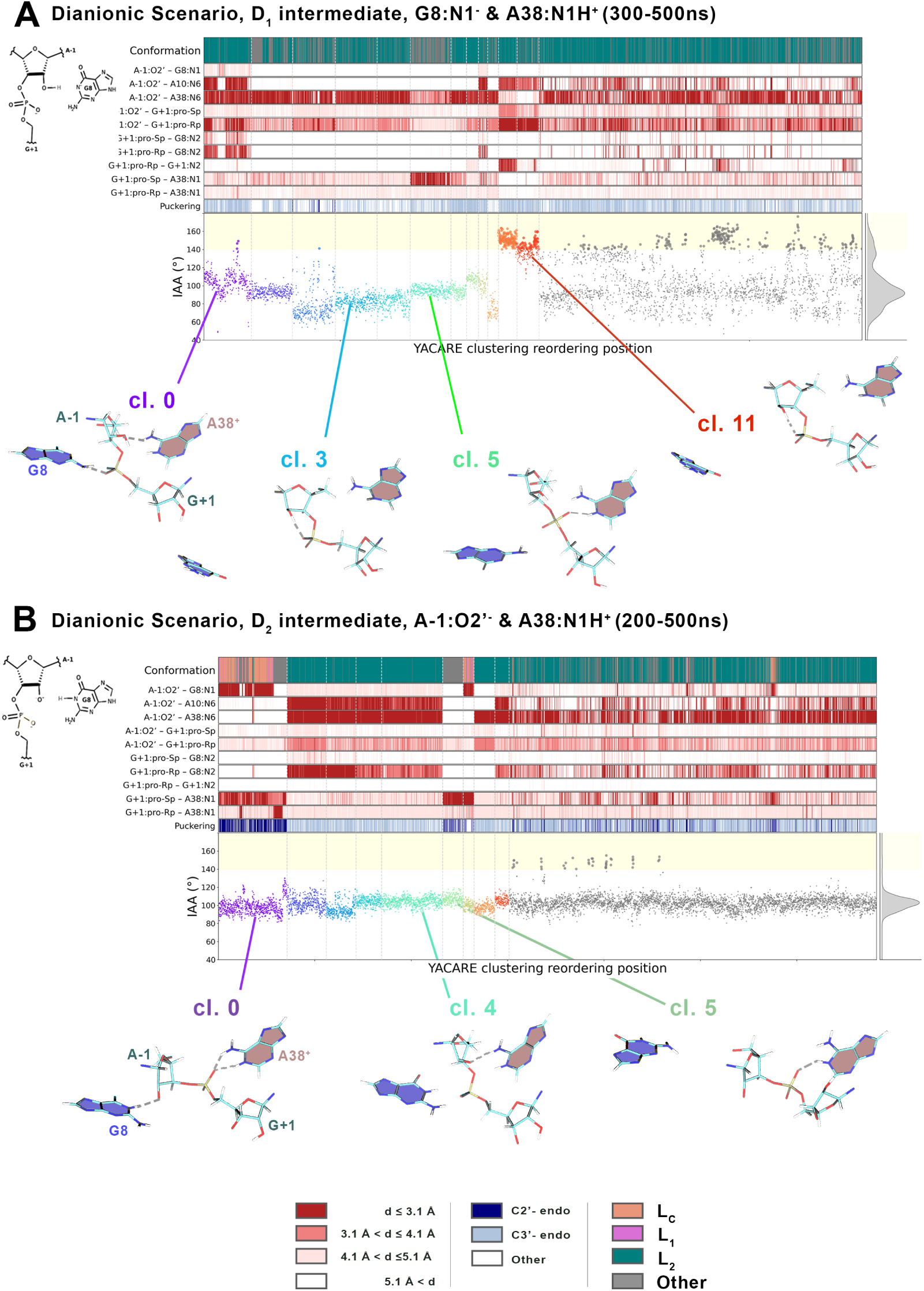
Evolution of key catalytic site observables for dianionic intermediates, with frames ordered by clustering analysis: (A) Deprotonation of the G8*^−^* D_1_ intermediate and (B) Activation of O2’*^−^* by the basic G8 D_2_ intermediate (see Figure 2). Row 1: conformational label (see Methods and Figure 3 for conformational labeling strategy); rows 2–10: heatmap of the key catalytic site distances; row 11: A-1 puckering pseudo-rotation conformations; row 12: inline attack angle (IAA) values. Below: representative structures from major clusters with hydrogen bonds (represented by dashed lines).

**Figure S5:**
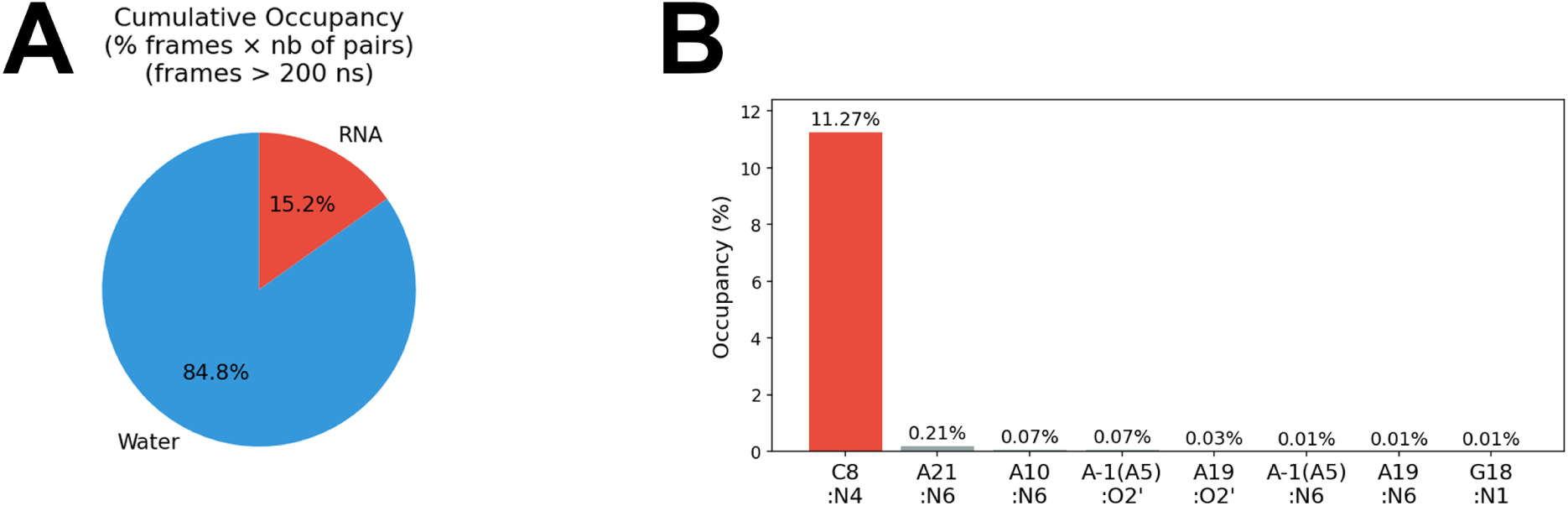
Hydrogen bond partners of G8*^−^* (N1*· · ·* H-X) in the D_1_ protonation state, computed over the converged portion of the REST2 simulation (200-500ns, see Figure S1 D). (A) Breakdown by partner type (solvent versus RNA). (B) Occupancy of individual RNA acceptors, expressed as a fraction of all hydrogen bonds formed.

**Figure S6:**
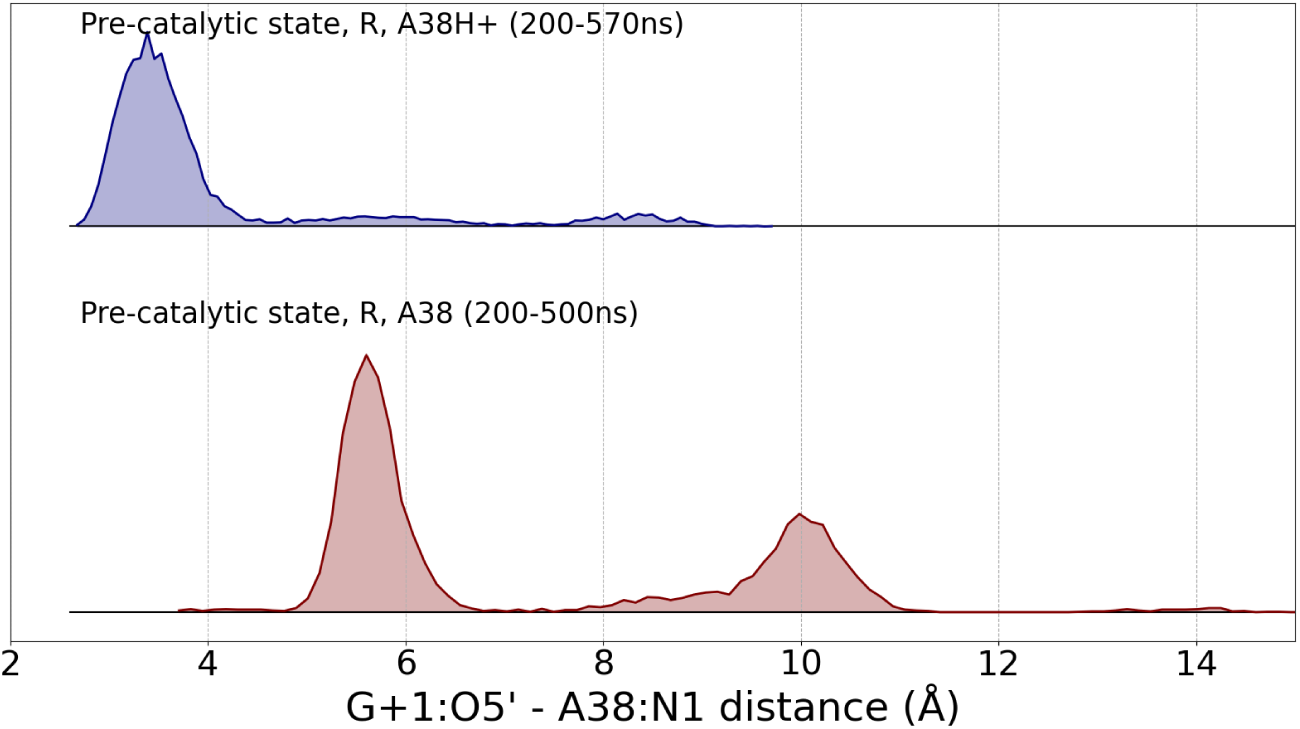
Distributions of the G+1:O5’*· · ·* A38:N1 distance from two REST2 simulations of the pre-catalytic state R: (top) A38 protonated at N1, as in all other systems of this study (see Figure S1 A); (bottom) A38 deprotonated at N1. Convergence times are indicated in parentheses.

